# Inter-basin water transfers and the expansion of aquatic invasive species

**DOI:** 10.1101/356923

**Authors:** Belinda Gallardo, David C. Aldridge

## Abstract

Inter-basin Water Transfers (IBWT) are recognized as one of the major pathways of freshwater invasion. They provide a direct link between previously isolated catchments and may modify the habitat conditions of the receiving waters such that they become more favourable for the establishment of invasive species. Combined, IBWT and invasive species will intensify the stress upon native species and ecosystems. Using the Severn and Thames Rivers-two of the largest river systems in Great Britain—as a case study, here we assess the potential influence of IBWT on the expansion of invasive species and thus their impact on biodiversity conservation. The Thames Valley is subject to extensive water abstraction, and an increasing population means that supplemented flow from the River Severn is being considered. Multi-scale Suitability Models, based on climate and water chemistry respectively, provided novel evidence that there is serious risk for further spread of invasive species in the focus area, particularly of the quagga mussel, a recent invader of the Thames River. Native freshwater mussels are particularly vulnerable to changing environmental conditions, and may suffer the decrease in alkalinity and increase in sedimentation associated with an IBWT from the lower Severn to the upper Thames. Regional models suggest considerable overlap between the areas suitable for three vulnerable native freshwater mussels and the expansion of invasive species that negatively impact upon the native mussels. This study illustrates the use of novel spatially-explicit techniques to help managers make informed decisions about the risks associated with introducing aquatic invasive species under different engineering scenarios. Such information may be especially important under new legislation (e.g. EU Invasive Species Regulation No 1143/2014) which increases the responsibility of water managers to contain and not transfer invasive species into new locations.

## 1. Introduction

Climate change is currently prompting water-engineering responses in anticipation of greater frequency of floods and droughts that will not only threaten water security but also freshwater biodiversity (Vörösmarty et al. 2010). Inter-Basin Water Transfers (IBWT) modify the water flow, chemistry and temperature of receiving waters, affecting the composition, abundance, richness and distribution of aquatic communities (Bunn and Arthington 2002). In fact, IBWTs are recognized as one of the major pathways of freshwater invasion in Europe (Jażdżewski 1980, Leuven et al. 2009). First, IBWTs provide a direct link between previously isolated catchments (Gupta and van der Zaag 2008). For example, the once isolated rivers draining the Caspian and Black Seas became inter-connected through canal construction in the 20^th^ century, facilitating the spread of Ponto-Caspian invasive species into western Europe (e.g. Rhine-Danube-Main waterways, Leuven et al. 2009). Interconnectivity of waterways has consequently removed major biogeographic barriers, leading to a situation where distant rivers share a high proportion of their (invasive) flora and fauna (Galil et al. 2007). Further, IBWTs modify the habitat conditions of the receiving waters, changes that often favour the establishment of invasive species (Dudgeon et al. 2006). For instance, increased alkalinity and temperature are known to provide a competitive advantage to invasive species over their native counterparts (Gallardo and Aldridge 2013c, Grabowski et al. 2009, Paillex et al. 2017). The compounded deleterious effects of water transfers and invasive species will intensify the stress upon native species, with the potential to ultimately provoke local extinctions (O’keeffe and De Moor 1988). For instance, the Tajo-Segura IBWT (SW Spain) operating since 1978, promoted the invasion of at least 13 invasive fish and one mollusc (Oliva-Paterna et al. 2014, Zamora-Marin et al. 2018), which resulted in 40% of native species threatened with local extinction (Oliva-Paterna et al. 2014).

To illustrate the potential influence of IBWTs on the expansion of invasive species and thus their impact on biodiversity conservation, here we use the catchments of the River Severn and River Thames as a case study (Fig. 1). Representing the two largest rivers in Great Britain, they collectively supply drinking water to 35 million people (>50% GB population). Climate simulations in Great Britain predict substantial reductions in summer precipitation accompanied by increased evapotranspiration throughout the year, leading to reduced flows in the Thames River in late summer and autumn (Diaz-Nieto and Wilby 2005). Water transfers from the nearby Severn River have long been suggested as an adaptation strategy to ensure the water supply of the Thames region (Jamieson and Fedra 1996, Rodda 2006). The Severn-Thames IBWT could arguably modify the river flow, hydrodynamics and water quality conditions of both rivers, the conservation of freshwater and terrestrial protected sites near the abstraction and discharge points as well as along the transfer route, and the movement of migratory fish (Bunn and Arthington 2002). Furthermore, the River Thames is one of the greatest hotspots of aquatic invasive species in the world (Jackson and Grey 2013, Zieritz et al. 2014), including myriad Ponto Caspian invaders with a high potential to spread widely across Great Britain (Gallardo and Aldridge 2015). Consequently, there is a serious risk of an eventual IBWT promoting further expansion of invasive species in the region, which may severely affect the status of vulnerable native biota and the broader ecosystem services of the two rivers. Of particular concern is the depressed river mussel (*Pseudanodonta complanata* Rossmassler, 1835) that has disappeared from over 30% of its historical sites in Great Britain (Killeen et al. 2004), the duck mussel (*Anodonta anatina* Linnaeus, 1758), and the painter’s mussel (*Unio pictorum* Linnaeus, 1758). The conservation of all three molluscs is a concern throughout Europe (Lopes-Lima et al. 2017). They are especially vulnerable to fouling from invasive zebra mussels (*Dreissena polymorpha* Pallas 1771) and quagga mussels (*Dreissena rostriformis bugensis* Andrusov 1897), and competition for habitat and resources with Asian clams (*Corbicula fluminea* O. F. Müller, 1774), all of which are present within the region (Aldridge et al., 2004; Sousa et al, 2011).

**Figure 1.**
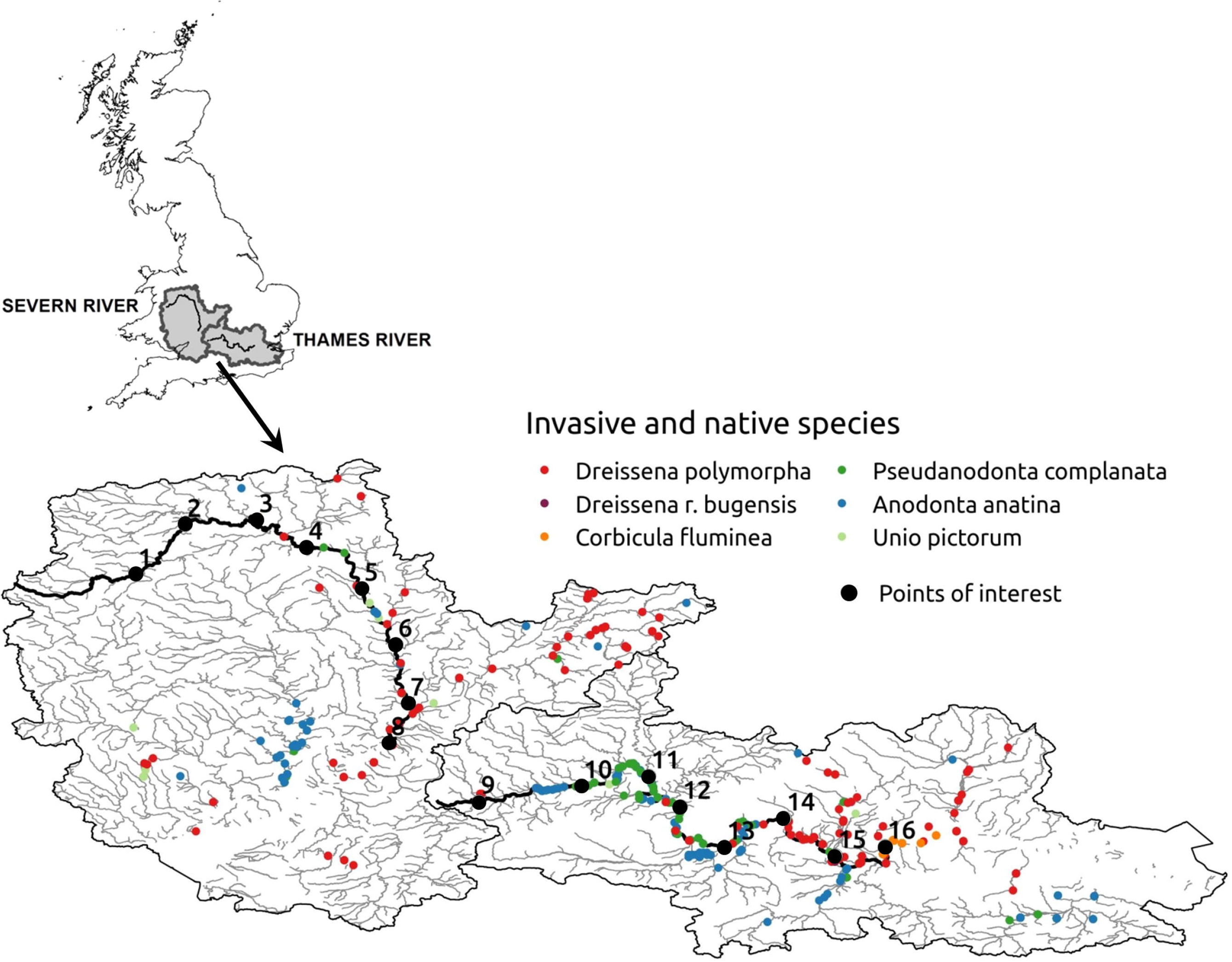
Study area located in the Severn and Thames river catchments in the south of Great Britain. The presence of three invasive and three native species is shown. Points of interest, located at regular 30-km intervals in each river, represent potential withdrawal and discharge river sections for an inter-basin water transfer.

This study investigates: i) the climate suitability for three high-risk aquatic invasive species, ii) the most likely pathways of spread between catchments, iii) the environmental suitability (in terms of river geomorphological and physicochemical characteristics) of the focus area to invasive species, and iv) their combined effects upon the conservation of three native species. Ultimately, this study evaluates the collective influence of water transfers on the potential for establishment and spread of invasive species, and their consequences for the conservation of vulnerable freshwater taxa. This information is critical to inform river management in a global change context, particularly under new legislation (e.g. EU Invasive Species Regulation No 1143/2014) that increases the responsibility of water managers to contain and avoid transferring invasive species into new locations.

## 2. Materials and Methods

The objectives of this study were achieved using multi-scale Species Distribution Models (SDM). These models are increasingly used to anticipate biosecurity threats associated with climate change and the expansion of invasive species (Jeschke and Strayer 2008). In brief, SDMs correlate the occurrence of a given species with the environmental conditions, usually climatic, of the sites it inhabits in order to locate areas that are most susceptible to invasion (Guisan and Thuiller 2005). While direct indicators of water conditions (water temperature and river flow) are preferable over indirect climate surrogates (air temperature and precipitation), multiple studies have demonstrated the prominent role of climate over geological (Quinn et al. 2014), hydrological (McGarvey et al. 2017) or socio-economic aspects (Gallardo and Aldridge 2013a) on the distribution of aquatic organisms at the global scale. Climate surrogates are especially useful when modelling invasive species because they allow using information from the entire global range of distribution of invasive species (native and invasive). This is important, because invasive species are rarely in equilibrium with the environment in the invaded range, and using regional information only may seriously underestimate their probability of further expansion. Many examples exist of studies using climate to model the potential distribution of aquatic organisms such as fish (e.g. Chen et al. 2007), aquatic snails (e.g. Loo et al. 2007), shrimps (e.g. Gallardo et al. 2012), crayfish (e.g. Capinha et al. 2012) and mussels (e.g. Drake and Bossenbroek 2004), among others. In addition, climatic factors offer the advantage of future projections that allow exploring the potential trajectories of invasive species, scenarios provided by the IPCC that are not yet available for other environmental or human indicators.

Nevertheless, we are aware that beyond temperature, local factors such as water chemistry and habitat structure are critical to narrow down the water bodies most likely to be invaded (Gallardo and Aldridge 2013c). For this reason, recent studies advocate for the integration of large-scale climate and regional-scale habitat conditions in a two-step modelling approach that makes use of all the available information to investigate the potential distribution of invasive species (Fournier et al. 2017, Gallardo and Aldridge 2013c, Gallien et al. 2012). Following this multi-scale approach, first, we use continental SDMs to locate broad areas suitable for the establishment of invasive species in the study area. We then evaluate the preferential corridors between both catchments that could eventually facilitate the expansion of invasive species. Finally, we use regional models to consider local-scale habitat conditions that ultimately determine the species survival, growth, spread and impact. By overlapping suitability maps for invasive and native species, we further identify water bodies where conflicts between native and invasive species are most likely to arise. Outputs from continental and regional SDMs are complementary and together will help evaluate biosecurity issues emerging from the IBWT.

### 2.1 Continental Species Distribution Models

Information on the current global (native and invaded) spatial distribution of the three invasive (*C. fluminea, D. polymorpha* and *D. r. bugensis*) and three native (*P. complanata, A. anatina* and *U. pictorum*) species was obtained from the following international and regional data gateways: Global Biodiversity Information Facility (GBIF, http://data.gbif.org), the National Biodiversity Network (NBN, Gateway http://data.nbn.org.uk), Discover Life (http://www.discoverlife.org), and complemented with an extensive ISI Web of Knowledge literature review (see list of references in Gallardo and Aldridge 2013c, Gallardo and Aldridge 2015). The zebra mussel, *D. polymorpha*, is widely spread across both catchments, whereas *D. r. bugensis* and *C. fluminea* are limited to the lower Thames River (Fig. 1). Native species are currently concentrated in the middle and upper sections of the Thames River and some of its tributaries, and in the middle reaches of the Severn River (Fig 1). Global maps for invasive species can be consulted in Supplementary Materials (Fig. SI).

To investigate the climatic suitability of the focus area to invasive and native species, five global bioclimatic variables plus geographic elevation (m) were obtained from WorldClim (http://www.worldclim.org): maximum temperature of the warmest month (°C), minimum temperature of the coldest month (°C), precipitation of the wettest month (mm), precipitation of the driest month (mm), and precipitation seasonality (coefficient of variation) (mm). Variables were chosen to show correlation r<|0.7|, a well-accepted limit for SDM (Gallardo et al. 2017). These bioclimatic variables represent annual trends, seasonality and extremes that are appropriate to explain species survival (Hijmans and Graham 2006) and were thus selected based on their relevance to explain the large scale distribution of species (see Quinn et al. 2014). For example, air temperature is directly related to water temperature (Stefan and Preud’homme 1993), which affects the reproduction, growth, dispersal, metabolism and oxygen consumption of aquatic organisms (Griebeler and Seitz 2007, Jacobsen et al. 1997). Rainfall patterns affect the discharge and depth of rivers and lakes and therefore the likelihood of droughts and floods that can have marked effects on freshwater organisms (Nickus et al. 2010), although we expect precipitation-related variables to be less important than temperature in the models. We included altitude in our models because the associated high currents and low organic matter can limit the distribution of aquatic molluscs, regardless of temperature (Quinn et al. 2014). For instance, zebra mussels in Europe are rarely found above 500 m (Strayer 1991). We chose a 30 arc-second (approximately 1×1 km at the equator) resolution for variables used as predictors in SDM, a high resolution that allows the precise characterization of the aquatic species’ climate niche (Gallardo et al. 2015). To calibrate SDMs, we used the MaxEnt algorithm, which typically outperforms other methods based on predictive accuracy (Merow et al. 2013). For input, MaxEnt models use the dataset of species occurrences and the set of climatic predictors that might affect the likelihood of species establishment. The following modelling parameters were implemented in software MaxEnt v3.3k: convergence threshold = 100, maximum iterations = 500, prevalence = 0.5, 10,000 random pseudo-absences.

To compensate for any potential sampling bias in the species current distribution, we used the Global Accessibility Map produced by the European Commission (http://bioval.irc.ec.europa.eu/)(Nelson 2008), which measures the travel time needed to access from any pixel to the closest major city (i.e. >50,000 inhabitants). This bias indicator outperformed other methods to correct the initial sampling bias of distribution models (Fourcade et al. 2014), and has been used to compensate sampling bias for the modelling of freshwater invasive species in Europe (Gallardo et al. 2017).

A 10-fold cross-validation was used to evaluate the predictive power of the model. This technique splits the occurrence dataset into ten equal-size groups called “folds”, and models are created leaving out each fold in turn. The omitted fold is then used for evaluation. Average values from the ten replicates were used for reporting and mapping potential distributions. To assess model performance, the Area Under the Receiving Operating Characteristic (ROC) Curve (AUC) (Hanley and McNeil 1982) was used, which represents the probability that a random occurrence locality is classified as more suitable than a random pseudo-absence. A model that performs no better than random has an AUC of 0.5, whereas a model with perfect discrimination scores 1.

After calibration, models were projected into the focus area to obtain suitability maps. Suitability is a measure of the match with the climatic conditions of locations currently invaded by a species and ranges from 0 (completely dissimilar) to 1 (perfect match). The threshold maximizing the sensitivity (i.e. number of presences correctly predicted) and specificity (i.e. number of absences correctly predicted) of the model was used to transform suitability maps into binary presence/absence maps. Binary maps allow investigating the full potential of spread of invasive species. By overlapping maps of native and invasive species, we identified regions where conflicts may arise.

### 2.2 Landscape Connectivity Analysis

Once suitability maps were obtained for each invasive species, we assessed the most likely natural pathways of dispersal between the two catchments using Landscape Analysis of Connectivity (LAC), a technique commonly used for examining population connectivity across landscapes (Urban et al. 2009). LAC assumes that high habitat suitability implies low cost to the dispersal for a species across a particular region, whereas regions with low suitability have high dispersal costs. Thus, LAC incorporates detailed habitat information as well as species-specific aspects on a measure of connectivity (Adriaensen et al. 2003). This type of modelling tool is receiving growing attention in applied land and species management projects as a powerful approach for predicting population connectivity.

As input, LAC takes a friction layer and a set of “points of interest”, that is, the potential source and sink locations of propagules. In the absence of specific points of withdrawal and discharge, we manually located 16 points of interest regularly at 30-km distance along the River Severn (N=8) and the River Thames (N=8) (Fig. 1). The inverse of each climate suitability map was used as a friction layer, assuming that the lower the climatic suitability for a particular invader, the higher the habitat resistance or “cost” to its dispersal.

The result of LAC is a map that indicates the routes that would ultimately facilitate the dispersal of invasive species between the eight points of interest located in each catchment and that should thus be avoided by an eventual IBWT. However, we must note that LAC is pertinent for assessing the risk of open-air canals, but irrelevant in cases where water is pumped through underground pipelines. While in this case the risk of invasive species transportation is mostly unknown, we will discuss the risks that our three focus invasive species may pose to the correct functioning of underground pipes. Friction layers and LAC were calculated using the SDMToolbox implemented in ArcView v.10.2.

### 2.3 Regional Species Distribution Models

The same set of invasive and native species was considered for modelling at the regional scale. The only exception was the quagga mussel, *D. r. bugensis*, since the information available for Great Britain (with only two presence points in the Wraysbury River, a tributary of the middle Thames) was insufficient to calibrate regional models.

Data on the current presence of both invasive and native species in Great Britain’s river networks used for continental models was complemented with information from the National Biodiversity Network (NBN) database (https://nbnatlas.org/). the UK Environment Agency (http://data.gov.uk/). Killeen (2012) and from our own field surveys in the upper Thames (data uploaded to NBN database and tagged to D. Aldridge).

Data on environmental predictors relevant to explain the regional distribution of species were obtained from the UK’s Environment Agency (EA) in the form of a shapefile. In this database, each segment in GB’s hydrological network is assigned an identification code by the EA, geographic coordinates, and a set of environmental values measured in 2012. The candidate environmental predictors comprised ten variables: alkalinity (ppm), river width (m), river depth (m), altitude (m), slope, substrate structure (% of boulders and pebbles, sand, silt and clay), discharge (m^3^/s), and ecological status (four categories: bad, moderate, good, not yet analysed) (see maps in Fig. S2). Such detailed river segment information is not available at larger continental to global scales, and for this reason could not be incorporated into continental models.

Regional distribution models were calibrated with MaxEnt v3.3k using the Samples With Data (SWD) option (Elith et al. 2010). Model settings (10-fold cross-replication), evaluation (by AUC) and mapping followed those described for continental SDM. Response curves provided by MaxEnt were used to investigate the influence of environmental variables likely to change after a water transfer such as alkalinity, discharge and the river substrate, on the establishment of invasive species and the survival of native species. Suitability maps for the three invasive and three native species were cross-compared to evaluate the potential for interaction between invasive and native species.

## 3. Results

Models calibrated with the global occurrence of invasive and native species and a set of climatic predictors showed a high accuracy (test AUC between 0.87 and 0.98, Table SI). The most important variables contributing to the models were minimum (48% on average) and maximum (22%) annual temperatures (Table S1).

Suitability maps presented in Figure 2 suggest a high potential for further spread of C. *fluminea* and *D. r. bugensis*, showing highest risk scores in the lower Thames River (Figs. 2C and 2E). While the zebra mussel, *D. polymorpha*, is widely established, models suggest it is likely to keep expanding in both catchments (Fig. 2A). According to the Landscape Analysis of Connectivity (LAC), the connection between the middle course of the Severn (point 5) and Thames (point 10) offers the most suitable corridor for the spread of *D. polymorpha* (Fig. 2B). The other two invaders, *C. fluminea* and *D. r. bugensis*, show preference for the main river channels and the connection between the upper Severn (point 7) and upper Thames (points 9 and 10) (Figs. 2 D and F).

**Figure 2.**
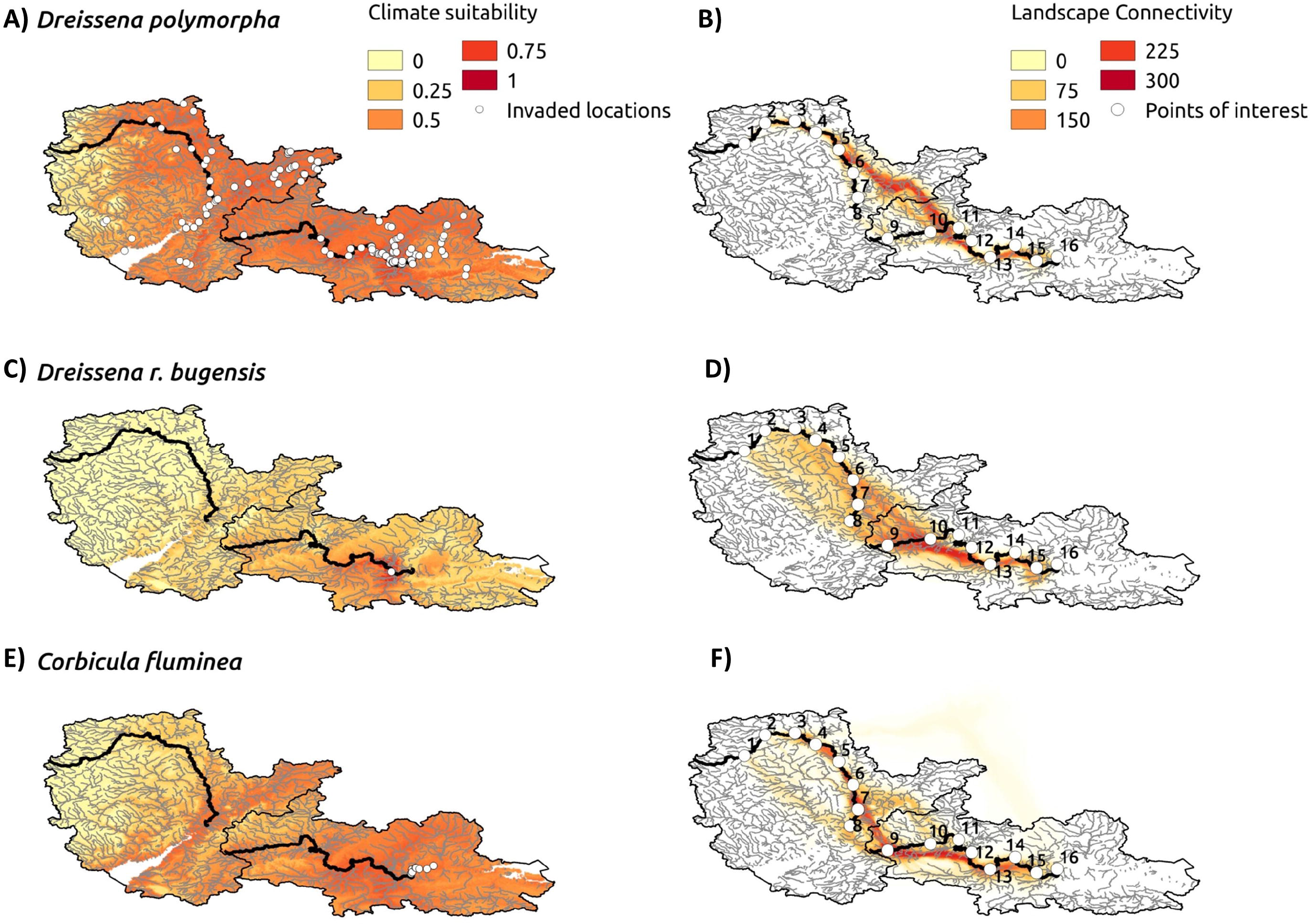
Climate suitability and landscape connectivity for three invasive species in the Severn and Thames river catchments. An inter-basin water transfer would increase propagule pressure, particularly in those areas with a high connectivity. Climate Suitability Models (A, C and E) indicate the probability of establishment if the species is introduced. White dots indicate locations already invaded by each species. Landscape Connectivity Analysis (B, D and F) indicates preferential connectivity pathways between potential withdrawal and discharge sections (points of interest) in both catchments.

Only the upper part of the Severn River (point 8) is suitable to the three invaders (Fig. 3A), whereas most of the potential discharge points in the Thames River (points 9-10 and 12-15) are highly vulnerable to invasion (Fig. 3A). Figure 3C shows potential areas of conflict with native species, which should be avoided by an IBWT to limit biosecurity threats. This map highlights the potential of the Severn catchment to offer refugia for the conservation of native molluscs (Fig. 3B), because the suitability for invasive species is concentrated in coastal areas and the lower valleys.

**Figure 3.**
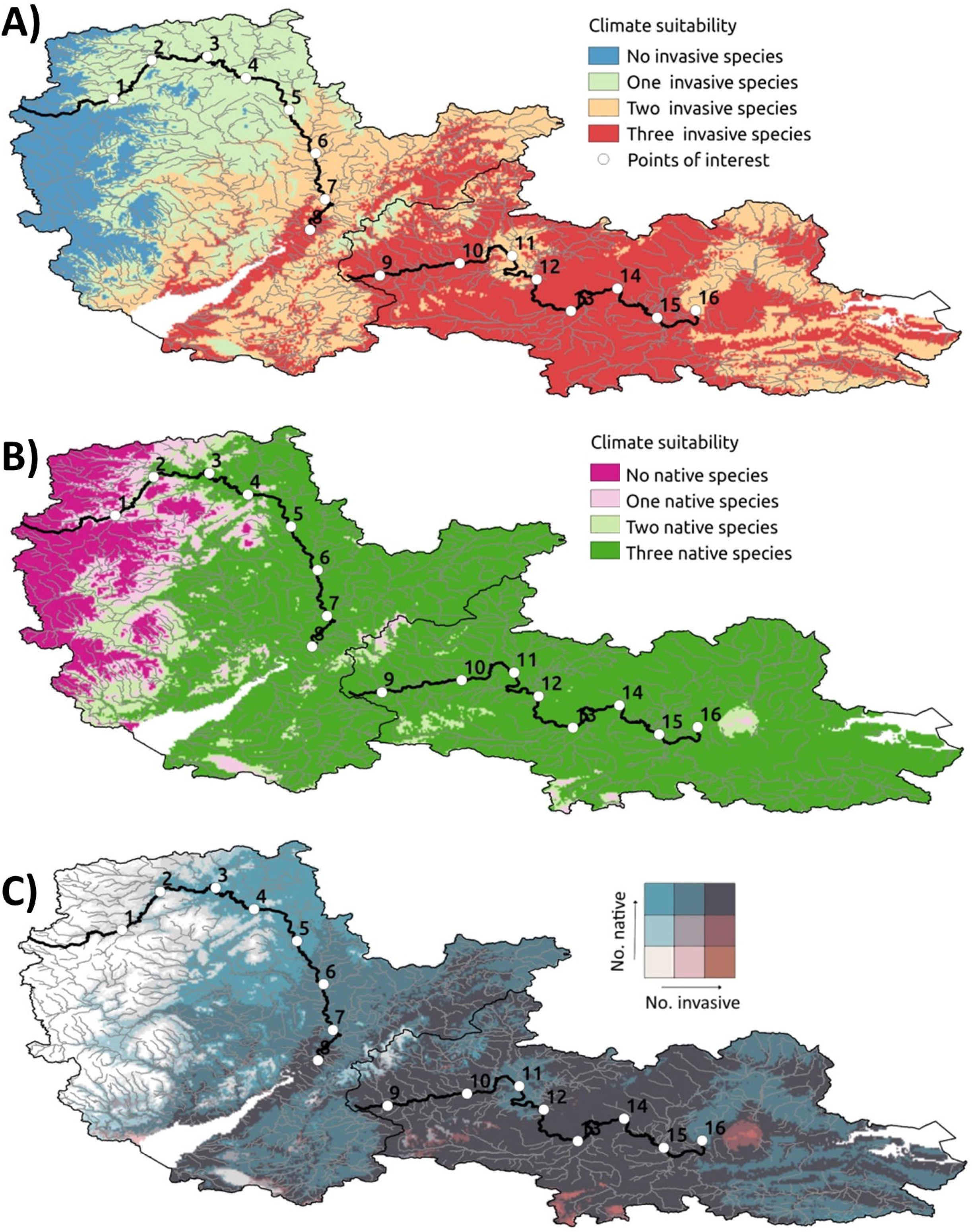
Cumulative climate suitability forthree invasive species (A), three native species (B) and their overlap (C) in the Severn and Thames river catchments. Points of interest, located at regular 30-km intervals in each river, represent potential withdrawal and discharge river sections for an inter-basin water transfer.

The geomorphological and physicochemical conditions of the Severn and Thames Rivers significantly differ (Table 1). The Severn waters have lower alkalinity and proportion of sand, but higher discharge than those of the Thames. Also, the ecological status of the Severn River is predominantly “Moderate” as opposed to “Poor” in the Thames (Table 1).

**Table 1.** River characteristics at 16 points of interest located regularly at 30-km distance in the Severn and the Thames Rivers (see Fig. 1). Results from Kruskal-Wallis performed between rivers are reported in the last row.

| River | Width<br>(m) | Depth<br>(cm) | Alkalinity<br>(ppm) | Silt<br>& clay<br>(%) | Sand<br>(%) | Pebbles<br>&<br>boulders<br>(%) | Discharge<br>(m<sup>3</sup>/s) | Ecological<br>Status |
| --- | --- | --- | --- | --- | --- | --- | --- | --- |
| 1-Thames | 55.30 | 260.00 | 182.00 | 4 | 5 | 90 | 40-80 | Poor |
| 2-Thames | 3.20 | 31.67 | 300.00 | 60 | 23 | 12 | <0.31 | Moderate |
| 3-Thames | 11.60 | 19.33 | 168.10 | 15 | 7 | 73 | 0.31-0.62 | Poor |
| 4-Thames | 3.20 | 31.67 | 300.00 | 60 | 23 | 12 | <0.31 | Moderate |
| 5-Thames | 1.40 | 8.00 | 240.00 | 50 | 47 | 3 | <0.31 | Moderate |
| 6-Thames | 5.80 | 46.67 | 169.00 | 8 | 75 | 15 | 1.25-2.5 | Poor |
| 7-Thames | 1.50 | 15.00 | 230.00 | 94 | 5 | 0 | <0.31 | Poor |
| 8-Thames | 1.70 | 8.67 | 215.00 | 17 | 43 | 40 | <0.31 | Moderate |
| 9-Severn | 30.00 | 50.00 | 105.27 | 100 | 0 | 0 | >80 | Moderate |
| 10-Severn | 26.70 | 233.33 | 119.10 | 78 | 8 | 13 | >80 | Moderate |
| 11-Severn | 50.00 | 300.00 | 178.00 | 100 | 0 | 0 | 40-80 | Moderate |
| 12-Severn | 38.70 | 25.00 | 95.80 | 2 | 0 | 37 | 40-80 | Moderate |
| 13-Severn | 30.70 | 24.33 | 18.10 | 0 | 7 | 33 | 10-20 | Moderate |
| 14-Severn | 7.70 | 14.00 | 120.10 | 8 | 17 | 42 | 0.62-1.25 | Moderate |
| 15-Severn | 8.50 | 121.67 | 59.80 | 77 | 0 | 20 | <0.31 | Poor |
| 16-Severn | 0.90 | 3.33 | 146.00 | 0 | 30 | 70 | <0.31 | Moderate |
| Kruskal-Wallis<br>Test (Chi, P) | 2.48,<br>n.s. | 0.54,<br>n.s. | 9.94,<br>0.001 | 0.00,<br>n.s. | 3.84,<br>0.04 | 0.10,<br>n.s. | 3.50, n.s. | 2.45, n.s. |

Regional species distribution models showed moderate accuracy scores (test AUC between 0.72 and 0.83, Table S2). Altitude, slope, ecological status and alkalinity were the most important factors explaining the distribution of invasive species; whereas substrate size (% pebbles, boulders and sand) and alkalinity determined the presence of native molluscs (Fig. 4A). Generally, invaders showed preference for lowland rivers, with Bad to Moderate ecological status, sand-dominated substrate (>20%) and high alkalinity (>120 mg/L) (Fig. S3). Similar to their invasive counterparts, native species showed preference for lowland rivers, with slow river discharges (<20 m^3^/s), gentle slope, high alkalinity (>200 mg/L) and high percentage of sand, silt and clay (Fig. S4). The regional heat-map on Figure 4B and 4C reflects the number of invasive species that could eventually find suitable environmental conditions and potential refuges for native species in each river segment of the Severn and Thames catchments.

**Figure 4.**
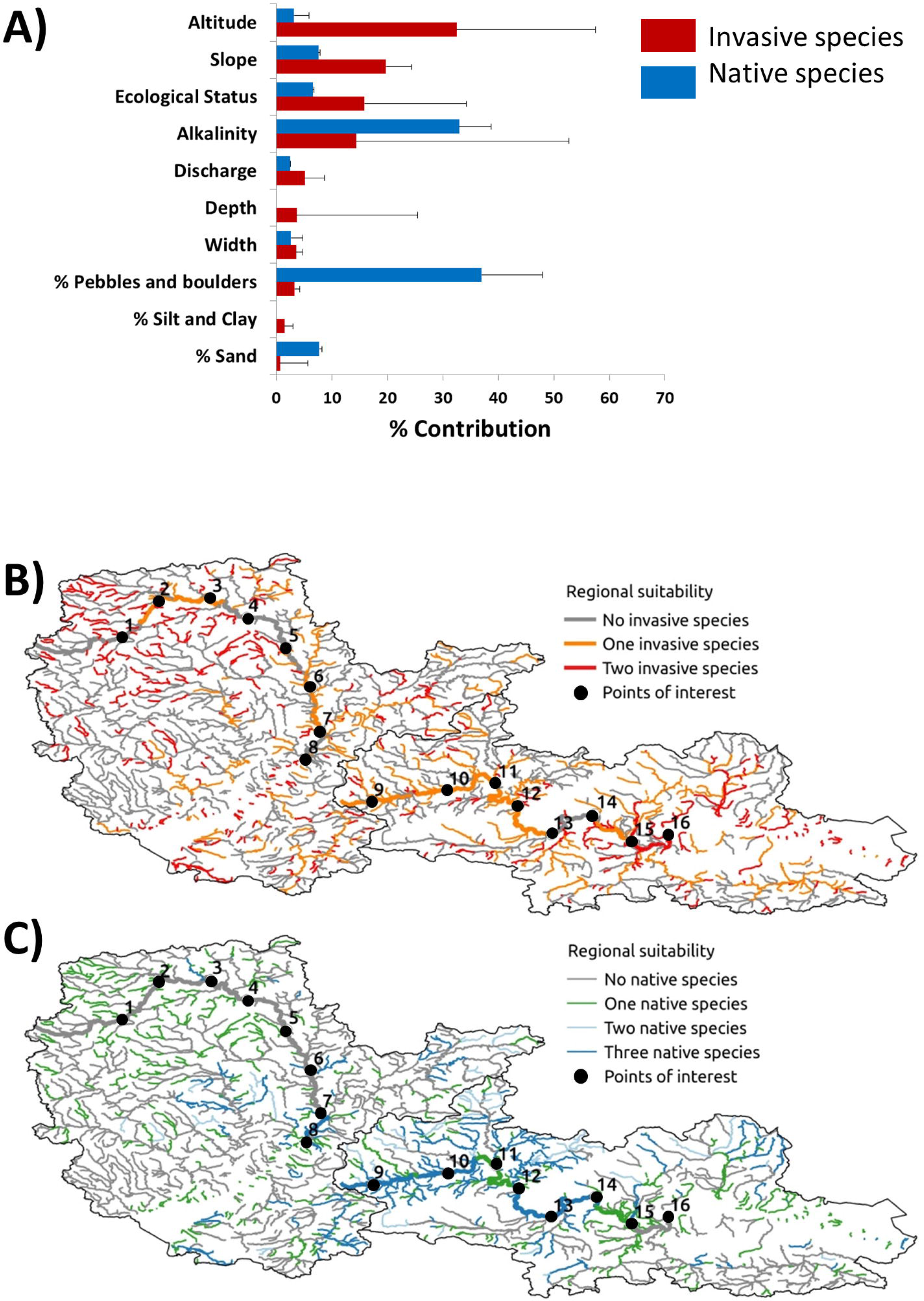
Results from regional species distribution models used to calculate habitat suitability for invasive and native species. A: Bars represent the mean and SD of the contribution of variables to regional models for invasive (red bars) and native (blue bars) species. B and C: cumulative regional suitability for two invasive species and three native species in the Severn and Thames river catchments. Points of interest, located at regular 30-km intervals in each river, represent potential withdrawal and discharge river sections for an inter-basin water transfer.

## 4. Discussion

### 4.1 Multi-scale risk assessment of invasive species

This study illustrates the use of species distribution models at multiple scales to evaluate the joint threat posed by an inter-basin water transfer and the concurrent expansion of invasive species upon the conservation of native freshwater vulnerable taxa. Multi-scale suitability models enabled us to assess the importance of different aspects of the process of invasion: establishment (climate and water chemistry suitability), spread (least-cost corridors) and impact (overlap with vulnerable native species).

### 4.2 Influence of climate on river invasibility

Based on climate suitability, the invasive species with the highest potential for spread in the study area is the quagga mussel, *D. r. bugensis*. Native from the Ponto Caspian region, the quagga mussel’s first British record was the Wraysbury River, a tributary of the middle Thames, in September 2014 (Aldridge et al. 2014), confirming previous risk assessments pointing to this mussel as the highest threat to British waters (Gallardo and Aldridge 2013d, Gallardo and Aldridge 2015, Roy et al. 2014). Latest investigations suggest that the species is spreading quickly, and that it may establish widely across England, west and southern Wales, and central Scotland (Gallardo and Aldridge 2015). According to our study, the suitability for this invader is highest in the middle-lower Thames and passive spread can allow rapid spread to downstream locations. For instance, Asian clams colonized the main channel of the Segura River (SW Spain) after the transfer with the Tajo River, and rapidly expanded downstream through passive drift (Zamora-Marín et al. 2018).

Most aquatic organisms are ectothermic and thus temperature is important in their physiology, bioenergetics, behaviour and biogeography (Jacobsen et al. 1997, Rahel and Olden 2008). Reflecting this, our models showed that minimum monthly temperature explained on average 48% of the distribution of invasive species, with low probability of establishment at values below freezing (0°C). For this reason, we may expect climate change to shift the climatic conditions of the focus area closer to the invaders’ optima. This may not only affect their probability of establishment, abundance, and distribution, but also their *per capita* effects (Hellmann et al. 2008, Rahel and Olden 2008), thereby increasing the stress upon native communities (lacarella et al. 2015).

### 4.3 Potential corridors between catchments

Using suitability as a friction layer, LAC allowed identification of the least-cost corridors between the lower Severn and upper Thames Rivers. A particularly high-risk route for a transfer would be an open-air canal from the upper Severn at point 7 to the upper Thames at point 9. This could result in the transfer of zebra mussels from the Severn to locations in the Thames where the species is not yet present. Furthermore, biosecurity threats are not limited to the points of withdrawal and discharge, but extend to the transfer route and engineering works involving machinery and materials moving between both rivers. The risk may be especially high in the upper reaches of the Thames that are inaccessible to boat traffic and rarely visited by anglers, and so at present the potential for human-mediated upstream introductions of invasive species are limited. Corridors may not only accelerate the expansion of invasive species, but also the genetic mixing of populations established in both rivers, particularly of the zebra mussel, thereby increasing their genetic heterogeneity and long-term adaptive potential (Facon et al. 2008). Genetic introgression from invasive carps has been already documented in the Segura River after the transfer from the Tajo (Oliva-Paterna et al. 2014)

The biosecurity risks posed by an eventual IBWT are high, but results from our LAC models only apply to open-air canals and not necessarily to the underground pipes that could alternatively connect the two rivers. Factors affecting the capacity of invaders to survive or even establish in underwater pipelines are not currently known. However, the three invasive species investigated here are important biofoulers known to affect irrigation systems (Rosa et al. 2011), and water treatment facilities (Elliott et al. 2005), causing multimillion losses in cleaning and maintenance (Oreska and Aldridge 2011). One of the main problems related to the presence of biofoulers is the clogging of pipelines, which reduces the water flow and often ceases irrigation. As a result, much more frequent cleaning is necessary. For instance, in the irrigation system fed by Mondego River (Portugal), several tonnes of Asian clams are removed from the canal in each of the cleaning routines taking place after the arrival of the species in 2005 (Rosa et al. 2011). To avoid such infestations, the Severn-Thames transfer scheme should plan effective measures to prevent the colonization of underground pipelines by biofoulers. This could include, whenever possible, operating transfers outside the breeding season of the mussels to prevent transfer of planktonic veliger larvae, or allowing pipelines to dry out between transfers so that biofouling communities cannot establish. Control of invasive bivalves within transfer systems could be achieved using approved, targeted, degradable technologies such as microencapsulated BioBullets (Aldridge et al. 2006).

In addition to the new connection and changes in habitat conditions, water transfers involve the construction of canals, embankments and reservoirs in both the donor and receiving catchments that secure hydrological stability during floods and droughts, thereby promoting water-related activities such boating, shipping or fishing that are important vectors for aquatic invasion (Carlton 1993). For instance, while 13 invasive fish were documented shortly after the Tajo-Segura connection was established, it is difficult to ascertain whether fish arrived through the new connection, or were intentionally introduced by fisherman in the associated reservoirs (Oliva-Paterna et al. 2014). An increase in propagule pressure, even in a populated catchment like the Thames, is not trivial, since this is the most consistent determinant of the establishment success of invasive species (Cassey et al. 2018). Furthermore, despite high habitat suitability along much of its length, invasive species are currently concentrated in the lower sections of the Thames River (except the zebra mussel, Fig. 2). This suggests that increased propagule pressure could facilitate further invasion into the upper-middle sections of the Thames and Severn Rivers.

### 4.4 Influence of water chemistry on river invasibility

At the regional scale, invaders showed preference for lowland rivers, with sandy substrate, and high alkalinity (>120 mg/L). Alkalinity reflects the availability of CaC0_3_ in water and therefore affects the formation of mussel shells (Greenaway 1985). For this reason, the quagga and zebra mussels colonize more frequently waters with relatively high alkalinity (Gallardo and Aldridge 2013c); a requirement that may explain the lower establishment of Ponto Caspian invaders in tributaries that usually register lower alkalinity than the main river channel (Grabowski et al. 2009). Remarkably, invasive species were more likely to occur in waterbodies with a Bad to Moderate ecological status, than in sites with a Good ecological status as defined by the UK Environment Agency, often based on invertebrate indicators. It is logical to expect a higher propagule pressure in waterbodies that are affected by various human activities and therefore show poor ecological status. Moreover, disturbance from human activities may especially favour the establishment of freshwater invasive species (Paillex et al. 2015). This observation could also suggest certain “biotic resistance” of well-preserved ecosystems to the colonization of invasive species, a process with mixed support in the literature (see Jeschke et al. 2012 for a review). If a relation between ecological status and either propagule pressure or actual invasion levels is confirmed, then promoting the ecological conservation of freshwaters may help prevent the spread of invaders. Conversely, sites with poor ecological status may be especially vulnerable through the facilitative interactions of invasive species which can lead to an “invasional meltdown” (Gallardo & Aldridge, 2015).

Potential changes in the Thames River water chemistry after an eventual IBWT include a decrease of alkalinity, and increase of siltation and discharge. While invasive species have preference for high alkalinity (Gallardo and Aldridge 2013c), the IBWT is unlikely to affect established populations of invasive species that are normally tolerant to a broad range of conditions. The three invasive species evaluated require > 20% silt and clay substrate to colonize, with decreasing suitability at increasing sand and boulders coverage (Fig. S3). Accumulative siltation after an IBWT would therefore favour the establishment of invasive species in the upper Thames, where the percentage of silt and clay tend to be low. The effect of increasing river discharge is difficult to anticipate. Some invaders such as the Asian clam show preference for low discharge (< 5 m^3^/s according to our database) and may be negatively affected by a potential increase in flow. Nonetheless, considering the low discharge of the Thames River (0.6-1.2 m^3^/s, Table 1), the water transfer is unlikely to affect the distribution of invaders in the study area.

Should an IBWT decrease light penetration to the river bed in the recipient region, such as through increased water depth or elevated turbidity, both quagga and zebra mussels might expect to benefit. Distributions of both species is strictly limited by UV light penetration which is fatal to their veliger larvae (Seaver et al. 2009), preventing the species from establishing in shallow, clear waters.

### 4.5 Collective impacts on native species

According to our maps, most of the focus area offers suitable environmental conditions to the establishment of the three native mussels (Figs. 3 and 4), and yet they are not usually found in the upper parts of rivers. This can be attributed primarily to the dietary requirements of unionid mussels that are collectors, requiring a relatively high level of particulate organic matter in suspension. Supplementation of flow in the upper Thames through an IBWT from the lower Severn may encourage native mussels to establish further upstream in the Thames. However, other aspects of such an IBWT could be harmful to the native mussels such as a decrease in alkalinity and increase in sedimentation, which is especially harmful to the depressed river mussel (Killeen et al., 2004). Furthermore, native freshwater mussels are particularly sensitive to river management operations (McIvor and Aldridge 2007), and may severely suffer the consequences of any water transfer between the Severn and the Thames Rivers.

In addition to the IBWT, climate warming may strongly affect the survival, growth and reproduction success of native mussels (Aldridge 1999, McIvor and Aldridge 2007). For instance in the Saone River (France), a 1.5°C increase in temperature resulted in a progressive change in freshwater mollusc communities, a decrease in species abundance, and richness (Mouthon and Daufresne 2010). Such changes in community structure usually lead to changes in important ecosystem functions such as nutrient cycling and benthic-pelagic coupling (Spooner and Vaughn 2008). Maximum temperature tolerance for the three native mussels are not known, but studies suggest that this limit is usually higher for invasive than native mussels, implying that global warming will disproportionally affect native species (Verbrugge et al. 2012). Further, differential effects of temperature in freshwater mussels and their fish hosts may cause problems by uncoupling the timing of mussels and fish reproduction as well aschanges in overall fish host availability (Hastie et al. 2003). This is not the case of invasive mussels such as the Asian clam or zebra and quagga mussels that reproduce via planktonic larvae, and this may further increase the competitiveness of invasive mussels under a climate change scenario (Gallardo and Aldridge 2013b).

There is little evidence about the impacts of the quagga mussel upon the UK’s native mussel populations, but we can expect them to be similar to the zebra mussels’ or even greater, since the quagga is able to colonize sites currently unreachable by the zebra mussel (Higgins and Zanden 2010). In particular, zebra mussels are known to impede the correct opening and closure of the valves of native mussels, to hamper their movement and burrowing, and to directly compete for space and resources (Karatayev et al. 1997). The expansion of the quagga mussel in the River Thames and its potential arrival to the Severn therefore represents a serious threat to the conservation of native freshwater mussels. Finally, engineering works associated to an eventual IBWT may exacerbate this problem (Mclvor and Aldridge 2007) and cause considerable habitat degradation and mussel mortality downstream of the discharge points in the Thames River.

### 4.6 Study limitations and applicability

Spatially-explicit techniques such as SDM and LAC have seen considerable application for land and species management, and yet they have been rarely used to model aquatic ecosystems. This is probably because aquatic information is often available as vector (point or segment) rather than raster (gridded) format. Here we have illustrated the application of spatially explicit techniques combining the available information at global and regional scales. Nevertheless, we must bear in mind that models have been calibrated using a (limited) number of predictors and that other missing factors (e.g. interaction with native species, microhabitat and food availability, barriers to dispersal, habitat disturbance) may affect the ultimate spread and successful establishment of invaders as well as the survival of native species. In addition, models reflect suitability, that is, probabilities of invasion in the event of an introduction, and not absolute survival limits. A high suitability does not necessarily mean the species will establish, but simply that conditions are suitable for this event. Since colonization of the upper section of rivers is often dispersal-limited, special attention should be paid to the introduction of propagules by means of engineering works associated with IBWT. Ultimately, this case study illustrates how SDM can broadly help anticipate the expansion of multiple potential invaders, thereby enabling informed prioritisation of limited resources to guide monitoring, management and control decisions.

## 5. Conclusions

- *Inter-Basin Water Transfers may promote the expansion of aquatic invasive species*. An IBWT can increase propagule pressure to recipient regions, change habitat conditions that may benefit invasive species and harm native species, and create habitats such as canals, embankments and reservoirs that promote water-related activities (boating, shipping or fishing) and that are important vectors for aquatic invasion.
- *The invasive species with the highest potential for spread in the Thames and Severn Rivers is the quagga mussel*, D. r. bugensis. The species is currently limited to one location in the Thames, but shows high climate suitability in the study area. Quagga mussels may be able to colonize sites currently unreachable by the zebra mussel and therefore represent a serious threat to the conservation of native freshwater mussels. Quagga and zebra mussels may establish further upstream if IBWT leads to increased water depth and higher turbidity.
- *Climate change may exacerbate the risks associated to an IBWT*. Climate change will shift the climatic conditions of the focus area closer to the invaders’ optima there by increasing their probability of establishment, abundance, and *per capita* effects upon native communities. At the same time, climate warming may reduce habitat suitability for native mussels and interfere with reproduction if temperature affects their fish hosts.
- *Native species will suffer the concurrent effects of habitat degradation, the arrival of invasive species and climate change*. Aspects of an IBWT harmful to the native mussels include a decline in alkalinity, increase in sedimentation, and river management operations that intensify the stress upon native mussels. The IBWT jeopardizes the potential role of the Severn River as refuge for the conservation of native mussels.
- *Management implications*. A particularly high-risk route for a transfer would be anopen-air canal from the upper Severn to the upper Thames. Since colonization of the upper section of both rivers is dispersal-limited, special attention should be paid to the introduction of propagules by means of engineering works that can cause considerable habitat degradation and native mussel mortality. The potential transfer of invasive species through IBWTs could be reduced by operating transfers outside their reproductive season so that planktonic larvae are not transported. Fouling of transfer systems may be reduced through operating the transfer sporadically thus allowing the pipeline to periodically run dry.

## 6. Authors contributions

BG and DCA designed the study. DCA acquired part of the data. BG conducted the analyses. BG and DCA wrote the manuscript.

## 7. Acknowledgements

BG was funded by a Juan de la Cierva fellowship by the Spanish Ministry of Economy (JCI-2012-11908). DCA was supported by a Dawson Lectureship from St. Catharine’s College, Cambridge. The main author is grateful for the technical assistance of Manuel Pizarro (IPE-CSIC), in this investigation.

## References

Adriaensen, F., Chardon, J., De Blust, G., Swinnen, E., Villalba, S., Gulinck, H. and Matthysen, E. (2003) The application of ‘least-cost’modelling as a functional landscape model. Landscape and urban planning 64(4), 233–247.

Aldridge, D.C. (1999) The morphology, growth and reproduction of Unionidae (Bivalvia) in a fenland waterway. Journal of Molluscan Studies 65, 47–60.

Aldridge, D.C., Elliott, P. and Moggridge, G. (2006) Microencapsulated BioBullets for the control of biofouling zebra mussels. Environmental science & technology 40(3), 975–979.

Aldridge, D.C., Ho, S. and Froufe, E. (2014) The Ponto-Caspian quagga mussel, Dreissena rostriformis bugensis (Andrusov, 1897), invades Great Britain. Aquatic Invasions 9(4), 529–535.

Bunn, S.E. and Arthington, A.H. (2002) Basic principles and ecological consequences of altered flow regimes for aquatic biodiversity. Environmental Management 30(4), 492–507.

Capinha, C., Anastácio, P. and Tenedório, J. (2012) Predicting the impact of climate change on the invasive decapods of the Iberian inland waters: an assessment of reliability. Biological Invasions 14(8), 1737–1751.

Carlton, J.T. (1993) Zebra mussels:biology, impacts, and control. Nalepa, T.F. and Schloesser, D.W. (eds), pp. 677–697, CRC Press, Boca Raton (Florida).

Cassey, P., Delean, S., Lockwood, J.L., Sadowski, J. and Blackburn, T.M. (2018) Dissecting the null model for biological invasions: A meta-analysis of the propagule pressure effect. PLOS Biology 16(4), e2005987.

Chen, P.F., Wiley, E.O. and Mcnyset, K.M. (2007) Ecological niche modeling as a predictive tool: silver and bighead carps in North America. Biological Invasions 9(1), 43–51.

Diaz-Nieto, J. and Wilby, R.L. (2005) A comparison of statistical downscaling and climate change factor methods: impacts on low flows in the River Thames, United Kingdom. Climatic Change 69(2), 245–268.

Drake, J.M. and Bossenbroek, J.M. (2004) The potential distribution of zebra mussels in the United States. Bioscience 54(10), 931–941.

Dudgeon, D., Arthington, A.H., Gessner, M.O., Kawabata, Z.I., Knowler, D.J., Lévëque, C., Naiman, R.J., Prieur-Richard, A.H., Soto, D. and Stiassny, M.L. (2006) Freshwater biodiversity: importance, threats, status and conservation challenges. Biological Reviews 81(2), 163–182.

Elith, J., Phillips, S.J., Hastie, T., Dudfk, M., Chee, Y.E. and Yates, C.J. (2010) A statistical explanation of MaxEnt for ecologists. Diversity and Distributions 17(1), 43–57.

Elliott, P., Aldridge, D., Moggridge, G.D. and Chipps, M. (2005) The increasing effects of zebra mussels on water installations in England. Water and Environment Journal 19(4), 367–375.

Facon, B., Pointier, J.P., Jarne, P., Sarda, V. and David, P. (2008) High genetic variance in life-history strategies within invasive populations by way of multiple introductions. Current Biology 18(5), 363–367.

Fourcade, Y., Engler, J.O., Rödder, D. and Secondi, J. (2014) Mapping species distributions with MAXENT using a geographically biased sample of presence data: A performance assessment of methods for correcting sampling bias. Plos One 9(5), e97122.

Fournier, A., Barbet-Massin, M., Rome, Q. and Courchamp, F. (2017) Predicting species distribution combining multi-scale drivers. Global Ecology and Conservation 12, 215–226.

Galil, B.S., Nehring, S. and Panov, V. (2007) Biological Invasions, pp. 59–74, Springer.

Gallardo, B. and Aldridge, D.C. (2013a) The ‘dirty dozen2019: socio-economic factors amplify the invasion potential of 12 high risk aquatic invasive species in Great Britain and Ireland. Journal of Applied Ecology 50(3), 757–766.

Gallardo, B. and Aldridge, D.C. (2013b) Evaluating the combined threat of climate change and biological invasions on endangered species. Biological Conservation 160, 225–233.

Gallardo, B. and Aldridge, D.C. (2013c) Priority setting for invasive species management: integrated risk assessment of multiple Ponto Caspian invasive species into Great Britain. Ecological Applications 23(2), 352–364.

Gallardo, B. and Aldridge, D.C. (2013d) Review of the ecological impact and invasion potential of Ponto Caspian invaders in Great Britain, p. 120, Cambridge (UK).

Gallardo, B. and Aldridge, D.C. (2015) Is Great Britain heading for a Ponto-Caspian invasional meltdown? Journal of Applied Ecology 52(1), 41–49.

Gallardo, B., Aldridge, D.C., González-Moreno, P., Pergl, J., Pizarro, M., Pyšek, P., Thuiller, W., Yesson, C. and Vilà, M. (2017) Protected areas offer refuge from invasive species spreading under climate change. Global Change Biology 23(12), 5331–5343.

Gallardo, B., Errea, M. and Aldridge, D.C. (2012) Application of bioclimatic models coupled with network analysis for risk assessment of the killer shrimp, Dikerogammarus villosus, in Great Britain. Biological Invasions 14, 1265–1278.

Gallardo, B., Zieritz, A. and Aldridge, D.C. (2015) The importance of the human footprint in shaping the global distribution of terrestrial, freshwater and marine invaders. Plos One 10(5), e0125801.

Gallien, L., Douzet, R., Pratte, S., Zimmermann, N.E. and Thuiller, W. (2012) Invasive species distribution models - how violating the equilibrium assumption can create new insights. Global Ecology and Biogeography 21(11), 1126–1136.

Grabowski, M., Bacela, K., Konopacka, A. and Jazdzewski, K. (2009) Salinity-related distribution of alien amphipods in rivers provides refugia for native species. Biological Invasions 11(9), 2107–2117.

Greenaway, P. (1985) Calcium balance and moulting in the crustacea. Biological Reviews 60(3), 425–454.

Griebeler, E.M. and Seitz, A. (2007) Effects of increasing temperatures on population dynamics of the zebra mussel, Dreissena polymorpha: implications from an individual-based model. Oecologia 151(3), 530–543.

Guisan, A. and Thuiller, W. (2005) Predicting species distribution: offering more than simple habitat models. Ecology letters 8(9), 993–1009.

Gupta, J. and van der Zaag, P. (2008) Interbasin water transfers and integrated water resources management: Where engineering, science and politics interlock. Physics and Chemistry of the Earth, Parts A/B/C 33(1), 28–40.

Hanley, J.A. and McNeil, B.J. (1982) The meaning and use of the Area Under a Receiver Operating Characteristic (ROC) curve. Radiology 1, 29–36.

Hastie, L.C., Cosgrove, P.J., Ellis, N. and Gaywood, M.J. (2003) The threat of climate change to freshwater pearl mussel populations. Ambio 32(1), 40–46.

Hellmann, J.J., Byers, J.E., Bierwagen, B.G. and Dukes, J.S. (2008) Five potential consequences of climate change for invasive species. Conservation Biology 22(3), 534–543.

Higgins, S.N. and Zanden, M.J.V. (2010) What a difference a species makes: a meta-analysis of dreissenid mussel impacts on freshwater ecosystems. Ecological Monographs 80(2), 179–196.

Hijmans, R.J. and Graham, C.H. (2006) The ability of climate envelope models to predict the effect of climate change on species distributions. Global Change Biology 12(12), 2272–2281.

lacarella, J.C., Dick, J.T.A., Alexander, M.E. and Ricciardi, A. (2015) Ecological impacts of invasive alien species along temperature gradients: testing the role of environmental matching. Ecological Applications 25(3), 706–716.

Jackson, M.C. and Grey, J. (2013) Accelerating rates of freshwater invasions in the catchment of the River Thames. Biological Invasions 15(5), 945–951.

Jacobsen, D., Schultz, R. and Encalada, A. (1997) Structure and diversity of stream invertebrate assemblages: the influence of temperature with altitude and latitude. Freshwater Biology 38(2), 247–261.

Jamieson, D. and Fedra, K. (1996) The ‘WaterWare’decision-support system for river-basin planning. 3. Example applications. Journal of Hydrology 177(3–4), 199–211.

Jazdzewski, K. (1980) Crustaceana. Supplement, pp. 84–107, BRILL.

Jeschke, J., Gómez Aparicio, L., Haider, S., Heger, T., Lortie, C., Pyšek, P. and Strayer, D. (2012) Support for major hypotheses in invasion biology is uneven and declining. NeoBiota 14(0), 120.

Jeschke, J.M. and Strayer, D.L. (2008) Usefulness of bioclimatic models for studying climate change and invasive species. Year in Ecology and Conservation Biology 2008 1134, 1–24.

Karatayev, A.Y., Burlakova, L.E. and Padilla, D.K. (1997) The effects of Dreissena polymorpha (Pallas) invasion on aquatic communities in eastern Europe. Journal of Shellfish Research 16(1), 187–203.

Killeen, I., Aldridge, D.C. and Oliver, P.G. (2004) Freshwater bivalves of Britain and Ireland, FSC Publications, Shresbury, UK.

Killeen, IJ. (2012) A survey for the depressed river mussel Pseudanodonta complanata and other freshwater (unionid) mussels in the River Thames, p. 32, Malacological services for the UK Environment Agency.

Leuven, R., van der Velde, G., Baijens, I., Snijders, J., van der Zwart, C., Lenders, H.J.R. and de Vaate, A.B. (2009) The River Rhine: a global highway for dispersal of aquatic invasive species. Biological Invasions 11(9), 1989–2008.

Loo, S.E., Mac Nally, R. and Lake, P.S. (2007) Forecasting New Zealand mudsnail invasion range: Model comparisons using native and invaded ranges. Ecological Applications 17(1), 181–189.

Lopes-Lima, M., Sousa, R., Geist, J., Aldridge, D.C., Araujo, R., Bergengren, J., Bespalaya, Y., Bódis, E., Burlakova, L. and Van Damme, D. (2017) Conservation status of freshwater mussels in Europe: state of the art and future challenges. Biological Reviews 92(1), 572–607.

McGarvey, D.J., Menon, M., Woods, T., Tassone, S., Reese, J., Vergamini, M. and Kellogg, E. (2017) On the use of climate covariates in aquatic species distribution models: are we at risk of throwing the baby out? Ecography 41(4), 695–712.

Mclvor, A.L. and Aldridge, D.C. (2007) The reproductive biology of the depressed river mussel, Pseudanodonta complanata (Bivalvia: Unionidae), with implications for its conservation Journal of Molluscan Studies 73, 259–266.

Merow, C., Smith, M J. and Silander, J.A. (2013) A practical guide to MaxEnt for modeling species’ distributions: what it does, and why inputs and settings matter. Ecography 36(10), 1058–1069.

Mouthon, J. and Daufresne, M. (2010) Long-term changes in mollusc communities of the Ognon river (France) over a 30-year period. Fundamental and Applied Limnology/Archiv für Hydrobiologie 178(1), 67–79.

Nelson, A. (2008) Travel time to major cities: a global map of accessibility. Global Environment Monitoring Unit-Joint Research Centre of the European Commission, http://forobs.irc.ec.europa.eu/products/gam/.

Nickus, U., Bishop, K., Erlandsson, M., Evans, C.D., Forsius, M., Laudon, H., Livingstone, D.M., Monteith, D. and RThies, H. (2010) Climate change impacts on freshwater ecosystems. Kernan, M., Battarbee, R.W. and Moss, B. (eds), pp. 38–64, Blackwell Publishing Ltd., Oxford, UK.

O’keeffe, J. and De Moor, F. (1988) Changes in the physico-chemistry and benthic invertebrates of the great fish river, South Africa, following an interbasin transfer of water. Regulated Rivers: Research & Management 2(1), 39–55.

Oliva-Paterna, F.J., Verdiell-Cubedo, D., Ruiz-Navarro, A. and Torralva, M. (2014) La ictiofauna continental de la Cuenca del río Segura (SE Península Ibérica): décadas después de Mas (1986)/The freshwater ichthyofauna of the Segura river basin (SE Iberina Peninsula): decades after Mas (1986), p. 37, Servicio de Publicaciones, Universidad de Murcia.

Oreska, M. and Aldridge, D.C. (2011) Estimating the financial costs of freshwater invasive species in Great Britain: a standardized approach to invasive species costing. Biological Invasions 13(2), 305–319.

Paillex, A., Castella, E., zu Ermgassen, P., Gallardo, B. and Aldridge, D.C. (2017) Large river floodplain as a natural laboratory: non-native macroinvertebrates benefit from elevated temperatures. Ecosphere 8(10), e01972.

Paillex, A., Castella, E., zu Ermgassen, P.S.E. and Aldridge, D.C. (2015) Testing predictions of changes in alien and native macroinvertebrate communities and their interaction after the restoration of a large river floodplain (French Rhône). Freshwater Biology 60(6), 1162–1175.

Quinn, A., Gallardo, B. and Aldridge, D.C. (2014) Quantifying the ecological niche overlap between two interacting invasive species: the zebra mussel (Dreissena polymorpha) and the quagga mussel (D. rostriformis bugensis). Aquatic Conservation: Marine and Freshwater Ecosystems 24, 324–337.

Rahel, F.J. and Olden, J.D. (2008) Assessing the effects of climate change on aquatic invasive species. Conservation Biology 22(3), 521–533.

Rodda, J.C. (2006) Sustaining water resources in South East England. Atmospheric Science Letters 7(3), 75–77.

Rosa, I.C., Pereira, J.L., Gomes, J., Saraiva, P.M., Gonçalves, F. and Costa, R. (2011) The Asian clam Corbicula fluminea in the European freshwater-dependent industry: A latent threat or a friendly enemy? Ecological Economics 70(10), 1805–1813.

Roy, H., Peyton, J., Aldridge, D.C., Bantock, T., Blackburn, T., Bishop, J., Britton, R., Clark, P., Cook, E., Dehnen-Schmutz, K., Dines, T., Dobson, M., Edwards, F., Harrower, C., Harvey, M., Minchin, D., Newman, J., Noble, D., Parrott, D., Pocock, M., Preston, C., Roy, S., Salisbury, A., Schonrogge, K., Sewell, J., Shaw, R.E., Stebbing, P., Stewart, A. and Walker, K. (2014) Horizon-scanning for invasive alien species with the potential to threaten biodiversity in Great Britain. Global Change Biology 20(12), 3859–3871.

Seaver, R.W., Ferguson, G.W., Gehrmann, W.H. and Misamore, M.J. (2009) Effects of ultraviolet radiation on gametic function during fertilization in zebra mussels (Dreissena polymorpha). Journal of Shellfish Research 28(3), 625–633.

Spooner, D. and Vaughn, C. (2008) A trait-based approach to species’ roles in stream ecosystems: climate change, community structure, and material cycling. Oecologia 158(2), 307–317.

Stefan, H.G. and Preud’homme, E.B. (1993) Stream temperature estimation from air temperature. Journal of the American Water Resources Association 29(1), 27–45.

Strayer, D.L. (1991) Projected distribution of the zebra mussel, Dreissena polymorpha, in North America. Canadian Journal of Fisheries and Aquatic Sciences 48(8), 1389–1395.

Urban, D.L., Minor, E.S., Treml, E.A. and Schick, R.S. (2009) Graph models of habitat mosaics. Ecology letters 12(3), 260–273.

Verbrugge, L.H., Schipper, A., Huijbregts, M.J., Van der Velde, G. and Leuven, R.E.W. (2012) Sensitivity of native and non-native mollusc species to changing river water temperature and salinity. Biological Invasions 14(6), 1187–1199.

Vörösmarty, C.J., McIntyre, P.B., Gessner, M.O., Dudgeon, D., Prusevich, A., Green, P., Glidden, S., Bunn, S.E., Sullivan, C.A. and Reidy Liermann, C. (2010) Global threats to human water security and river biodiversity. Nature 467(7315), 555.

Zamora-Marín, J.M., Zamora-López, A., Sánchez-Pérez, A., Torralva, M. and Oliva-Paterna, F.J. (2018) Establecimiento de la almeja asiática Corbicula fluminea (Müller, 1774) en la cuenca del rfo Segura (SE Peninsula Ibérica). Limnetica 37(1), 1–7.

Zieritz, A., Gallardo, B. and Aldridge, D.C. (2014) Registry of non-native species in the Two Seas region countries (Great Britain, France, Belgium and the Netherlands). NeoBiota 23, 65–80.

